# Controlling for background genetic effects using polygenic scores improves the power of genome-wide association studies

**DOI:** 10.1101/2020.05.21.097691

**Authors:** Declan Bennett, Donal O’Shea, John Ferguson, Derek Morris, Cathal Seoighe

## Abstract

Ongoing increases in the size of human genotype and phenotype collections offer the promise of improved understanding of the genetics of complex diseases. In addition to the biological insights that can be gained from the nature of the variants that contribute to the genetic component of complex trait variability, these data bring forward the prospect of predicting complex traits and the risk of complex genetic diseases from genotype data. Here we show that advances in phenotype prediction can be applied to improve the power of genome-wide association studies. We demonstrate a simple and efficient method to model genetic background effects using polygenic scores derived from SNPs that are not on the same chromosome as the target SNP. Using simulated and real data we found that this can result in a substantial increase in the number of variants passing genome-wide significance thresholds. This increase in power to detect trait-associated variants also translates into an increase in the accuracy with which the resulting polygenic score predicts the phenotype from genotype data. Our results suggest that advances in methods for phenotype prediction can be exploited to improve the control of background genetic effects, leading to more accurate GWAS results and further improvements in phenotype prediction.

## Introduction

Linear mixed effects models (LMMs) are routinely applied to detect associations between SNPs and phenotypes in genome-wide association studies (GWAS) and many methods have been developed that enable these models to be applied efficiently to the large scale datasets that are typically now encountered in studies of complex traits^1–9^. Compared to fixed effects models for GWAS^10^, LMMs can be designed that have the advantage of being applicable to samples that include related individuals^1, 11, 12^. LMMs for this purpose typically include a random effect with covariance proportional to the kinship matrix that indicates the degree of relatedness between pairs of individuals in the sample^12^. The relatedness of individuals in the sample may be known *a priori* or may be derived from the genotype data by constructing a genetic relationship matrix (GRM), with entries corresponding to the genotypic covariance between pairs of individuals. When the entries of the GRM below a specified threshold are set to zero, the GRM is approximately equivalent to a family kinship matrix, with the degree of relatedness that the matrix captures controlled by this threshold. Thresholding the matrix to capture close family relationships (or cryptic relatedness^13^) allows specialized computational methods for sparse matrices to be applied so that model fitting remains tractable for studies that include large numbers of individuals^9^. This is the approach taken by fastGWA^9^, a recently developed tool that has been shown to generate correctly calibrated statistical results efficiently for biobank-scale GWAS.

In addition to enabling application to samples containing related individuals, LMMs can also account for genetic background effects^11, 14^. When a statistical model is used to test for a relationship between a given SNP (the test SNP) and a phenotype, the contribution of all genetic variants in the genome that are not in linkage disequilibrium with the test SNP is a form of background noise. If the trait of interest is both highly polygenic and highly heritable this noise may be substantial. Failure to account for sources of variance in the response in a statistical model can reduce the power to detect a relationship of interest^15, 16^. A LMM with a full GRM (i.e. derived from all SNPs in the data and with no threshold applied on the level of genetic correlation between individuals) is equivalent to a model in which all variants are assumed to have a causal effect on the phenotype, with effect sizes consisting of independent samples from a Gaussian distribution^8^. This is typically not a good fit to the true effect size distribution, and instead, the software package BOLT-LMM^8^ uses a spike-and-slab Gaussian mixture for the effect size distribution, with a component (the spike) close to zero corresponding to weak genome-wide effects and accounting for family relationships, and component with larger variance (the slab) corresponding to variants with large effects^8^. Fitting this more sophisticated model requires specialist numeric methods, that are relatively computationally intensive. Consequently BOLT-LMM is much more computationally intensive than fastGWA^9^.

The full GRM is an *N × N* matrix, where *N* is the number of individuals in the study. The memory and compute requirements of BOLT-LMM are kept tractable by selecting a subset of SNPs to include in the GRM, because BOLT-LMM performs operations that involve the GRM in a factorized form and requires *O*((*NM*)^1.5^) compute time, where *M* is the number of SNPs on which the GRM is based. Various options have been explored for which SNPs to include in the calculation of the GRM^11^. Including SNPs in LD with the target SNP results in loss of power, as the effect of the target SNP is partially accounted for by the random effect through the GRM. This has been referred to as proximal contamination^14^. On the other hand, including all (or most) SNPs that are not in LD with the target SNP, e.g. using a Leave One Chromosome Out (LOCO) approach, can result in dilution of the extent to which the relevant part of the genetic background is captured by the GRM. In the latter case, SNPs that are not relevant, in that they do not capture direct genetic effects or tag relevant population structure effects, effectively add noise to the GRM^14^. Alternatively, the GRM can be built from only the SNPs that are found using a linear model to be associated with the phenotype. Although this results in an increase in statistical power^11, 17, 18^, it does not fully control for population structure and is not recommended if population structure is of substantial concern^8, 11^. Methods have been developed that incorporate principal components into the GRM calculation built from significant SNPs; however, most of these methods are not suited to large biobank-scale data, without access to cloud computing or large compute farms^19–21^. Background genetic effects can also be included in the statistical model as fixed effects and this is the recommended approach when there are SNPs with large effect sizes^11^. A model fitting approach to determine the SNPs to include as fixed effects has been developed, and this also results in increased power in GWAS^14^.

As the genomic architecture of complex diseases is uncovered with the help of large biobanks, there is an advancing prospect of predicting quantitative phenotypes and the risk of complex diseases from genotype data. Recent years have seen substantial success and emerging clinical utility in phenotype prediction from polygenic scores (PGS)^22, 23^. PGS are constructed from weighted sums of allele dosages, with the weights corresponding to the effects size of the variants. Risk variants (variants associated with the phenotype) are typically inferred from the largest available GWAS, generally a meta-analysis. The clinical potential of PGS has already been shown in complex diseases such as coronary artery disease (CAD), diabetes and cancer^23–25^. In CAD, the identification of individuals with similar risk to those with rare high-risk monogenic variants has been reported^24^. Similarly, in breast cancer, pathogenic variants in BRCA1/2 account for 25% of familial risk of the disease with genome wide variants accounting for a further 18% of the risk^26, 27^. It is likely that in the future specialist machine learning methods will be developed to predict phenotype from genotype^23^, potentially achieving higher accuracy by incorporating the possibility of non-additive effects.

Here, we set out an approach to GWAS that seeks to separate the model fitting at the test locus and estimation of the genetic background effect. After carrying out an initial round of GWAS using an existing method, we derive a PGS for each chromosome, using the summary statistics for SNPs on the remaining chromosomes. This LOCO PGS is then included as a fixed effect in a second round of GWAS. We tested this approach in two ways. Firstly, using simulated data we tested for an improvement in power on the task of recovering known causal variants as a function of study size, number of causal variants and trait heritability. In addition, we applied the method to standing height data from the UK Biobank and determined the number and characteristics of additional variants that were detected. For an objective assessment of performance on real data, where the true associations are unknown, we divided the data into test and training sets and predicted the phenotype in the test set. The improvement in performance on the critical task of complex phenotype prediction illustrates the utility of the PGS as a means of accounting for off target genetic effects. This straightforward, modular approach to accounting for genetic background effects in GWAS has the advantage of leveraging advances in phenotype prediction as they become available. It also offers significant improvements in speed relative to existing methods that correct for genetic background.

## Results

We simulated data to evaluate the impact of including the LOCO PGS as a fixed effect in GWAS. The simulations consisted initially of a normally-distributed continuous trait in 100,000 individuals. The trait had a narrow-sense heritability (*h*^2^) of 0.5 and there were 1,000 causal SNPs with normally-distributed effects on the trait (see Methods for details). We incorporated the LOCO PGS as a fixed effect in a linear mixed model using GCTA fastGWA^9^ (we refer to this as fastGWA-PGS). To check the validity of this approach we performed simulations under the null model of no association between genotype and phenotype and found that the method was well calibrated (Fig. S1). This was in-line with our expectations as the LOCO PGS is approximately uncorrelated with the genotype of the tested SNP (see Supplementary Material for a mathematical justification). The p-values remained well calibrated even when we applied a higher p-value threshold for variants to include in the calculation of the LOCO PGS (Fig. S1), resulting in overfitting of the phenotype to SNPs not on the same chromosome as the test SNP. In 100 simulations we found that including a LOCO PGS resulted in a substantial improvement in power to detect the known causal SNPs (Fig. 1). We considered two alternative methods to select the SNP effects to include in the PGS calculation: pruning and thresholding (P&T) and LDpred2^28, 29^. When we included the PGS obtained using P&T as a fixed effect with fastGWA (which we refer to as fastGWA-PGS-PT) we recovered 82 additional causal variants, on average, below the conventional P-value threshold of 5*x*10^−8^ compared to fastGWA (corresponding to a relative increase in power of 18.4%; p = 3.0 × 10^−32^ from a paired T-test; Table S1-S3). The performance was further improved when we used LDpred2 to calculate the LOCO PGS (referred to as fastGWA-PGS-LDpred2). This resulted in the recovery of, on average, 115 more causal variants than fastGWA alone (relative increase of 25.9%; p = 2.3 × 10^−36^). We also simulated case control data for a binary traits with *h*^2^ of 0.5 and 1,000 causal loci, with disease prevalence, k, of 0.1 and 0.3. As with the quantitative trait simulations, inclusion of a fixed effect LOCO PGS always resulted in an increase in the average number of casual loci recovered, with an average of 28 more causal loci recovered for a disease prevalence of k=0.1 (p = 0.19) while, k = 0.3 recovered on average 48 more causal loci (p= 0.03) (Fig. S2 and Table S4 & S5).

**Figure 1.**
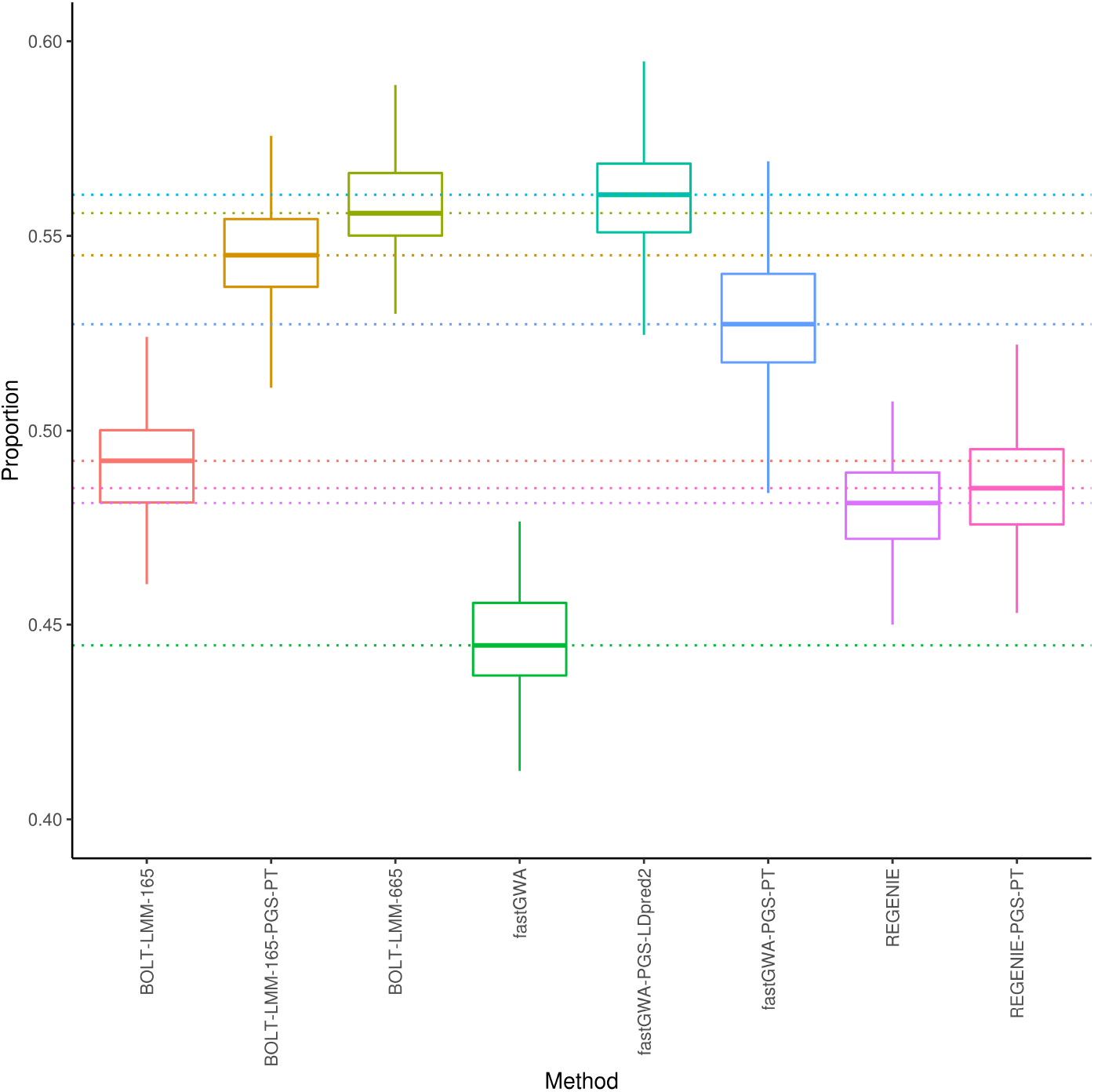
The proportion of causal variants recovered in 100 simulations. The boxplot shows the median (center line), upper and lower quartiles (hinges) and the maximum and minimum values not more than 1.5 times the interquartile range from the corresponding hinge (whiskers). The simulations consisted of 100,000 individuals and a continuous trait, with narrow-sense heritability of 0.5 and 1,000 causal variants.

The contribution to phenotype variance of background SNPs can also be modelled as a random effect in a linear mixed model. This approach is applied by BOLT-LMM, which uses a normal mixture random effect, with a component corresponding to SNPs with large effects. The running time of BOLT-LMM is proportional to *MN*^1.5^ and the memory requirement is approximately *MN*/4 bytes, where *N* is the number of individuals in the dataset and *M* is the number of SNPs included in the GRM^8^. When we ran BOLT-LMM with a subset of 165,683 SNPs (see Methods for how these were selected) we found that including the LOCO PGS as a fixed effect resulted in a substantial gain in power (Fig. 1), likely resulting from inability of the reduced GRM to account fully for genetic background. No further improvement was obtained by adding the LOCO PGS to BOLT-LMM with a GRM consisting of all of the 664,393 directly genotyped SNPs from the UKB (Fig. S3); however, the power obtained with the smaller GRM with the PGS fixed effect was close to the power obtained with the larger GRM, but with a much lower memory requirement (Table 1). Recently, a new fast method, REGENIE^30^, has been released that also includes control of the polygenic background effect based on prediction of the phenotype from SNPs that are not on the same chromosome as the test SNP. In our simulations the performance of REGENIE was intermediate between fastGWA and BOLT-LMM, but well behind fastGWA-PGS. REGENIE showed no improvement when the LOCO PGS was added as a fixed effect, suggesting that it accounts adequately for the genetic background effect. Note that REGENIE was omitted from Table 1, as the simulation is based on a single phenotype and would unfairly disadvantage REGENIE, which is optimized for the task of performing association analyses on multiple phenotypes simultaneously.

**Table 1.** Pipeline computation time and memory (N=100,000, M=665k)

| Method | CPU Time (s) |  |  |  | Max Memory (GB) |
| --- | --- | --- | --- | --- | --- |
|  | GWAS | LOCO PGS | GWAS(22 chr) | Total (CPU Time) |  |
| fastGWA-PGS-LDpred2 | 501.2 | 449.6 | 2,953.3 | 3,904.0 | 0.9 |
| fastGWA-PGS-PT | 501.2 | 245.8 | 2,953.3 | 3,700.2 | 0.7 |
| BOLT-LMM-665 | 119,202.0 | 0.0 | 0.0 | 119,202.0 | 15.5 |
| BOLT-LMM-165-PGS-PT | 92,108.0 | 245.8 | 614,514.4 | 706,868.2 | 3.9 |

We calculated receiver operator characteristic (ROC) curves to investigate whether the increased number of causal variants recovered when we included the LOCO PGS as a fixed effect reflected a reduction in P-values across the board for the phenotype-associated variants or also an improvement in the ordering of the variants, when the variants are ordered by the evidence of an association with the phenotype. Over 100 simulations we found that the area under the ROC curve (AUC) was always higher for fastGWA-PGS than for fastGWA without the LOCO PGS fixed effect. This was also the case for 99 of the 100 simulations when we added the PGS fixed effect to BOLT-LMM. The difference in sensitivity as a function of specificity (Fig. 2 and Table S6) showed that the sensitivity was consistently higher at a given specificity when the LOCO PGS-LDpred2 was included as a fixed effect, indicating an improvement in the ordering of the SNPs. The increase in mean sensitivity was up to 0.073 in the case of fastGWA-PGS-LDPred2 vs fastGWA, corresponding to a relative increase of 11.6% (at a specificity of 0.9988) over fastGWA. The addition of the LOCO PGS fixed effect led to a smaller but still consistent increase in sensitivity for BOLT-LMM-165. In this case, the greatest increase in the mean sensitivity was 0.028, corresponding to a 4.2% relative increase in sensitivity (at a specificity of 0.9991)

**Figure 2.**
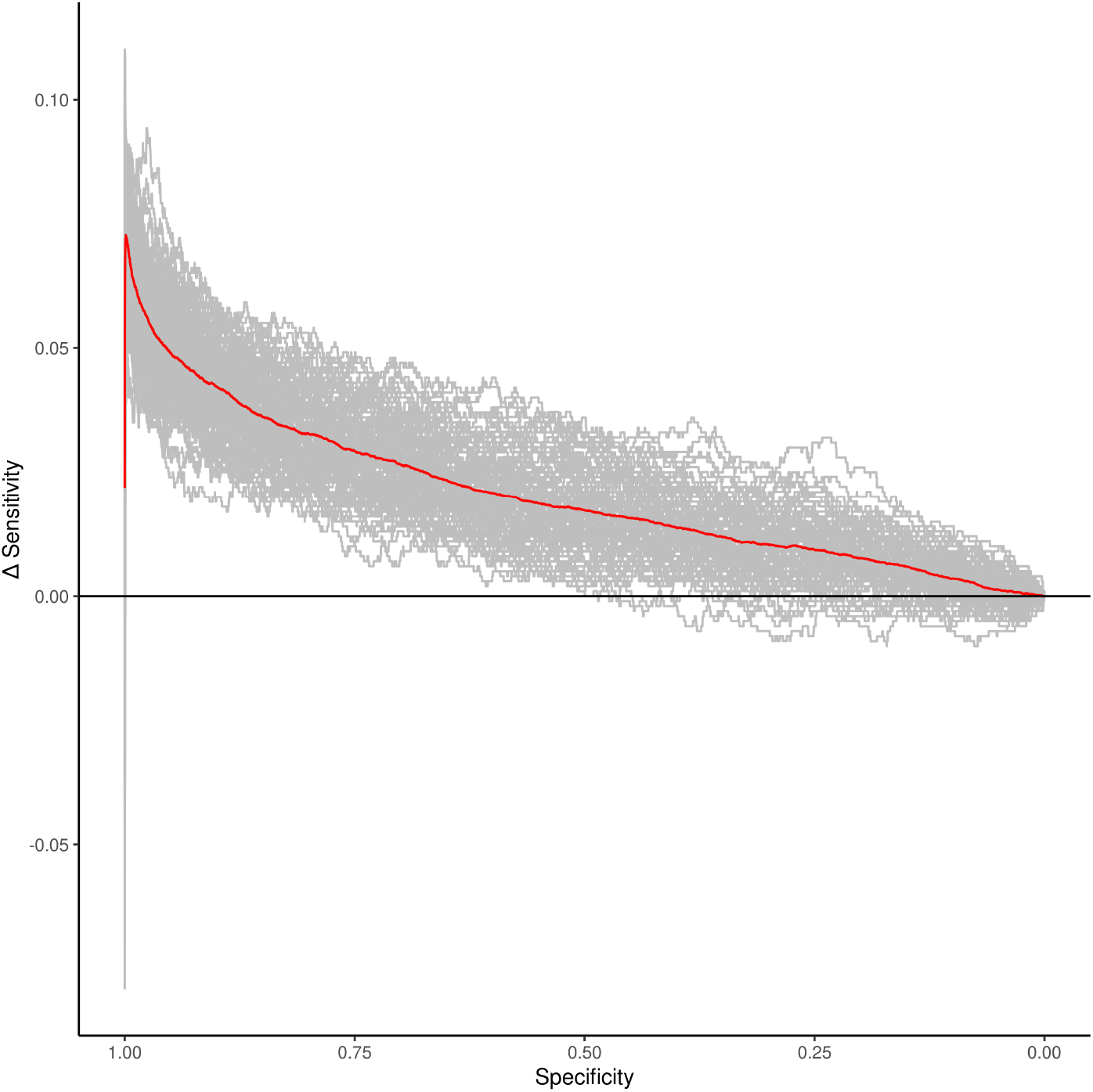
Difference in sensitivity (between fastGWA-PGS-LDpred2 and fastGWA) as a function of specificity for 100 simulations of a continuous trait with narrow-sense heritability of 0.5 and 1,000 causal variants in 100,000 individuals. The specificity (x-axis) is discretized in bins of size 0.0001. Each grey line shows the results of one simulation. The red line shows the mean difference over all simulations.

In addition to increasing the statistical power to detect causal variants, including the PGS fixed effect also resulted in an improvement in effect size estimates (Fig. S4). We found that when a fixed effect PGS was incorporated into the association study the median squared error (MEDSE) of the effect size estimate was substantially reduced (Fig. S4, Table S7-9). Interestingly, the MEDSE of the effect size estimate was largest across all methods for BOLT-LMM with the reduced GRM (Fig. S4).

### Effects of trait heritability, number of causal variants and sample size

We simulated data over a range of values of sample size, *h*^2^ and of the number of causal SNPs to investigate how these parameters affect the impact of including the LOCO PGS as a fixed effect on GWAS power. For the larger sample size, a small improvement in power was obtained even for the lowest values of *h*^2^ (0.1) simulated, with a statistically significant improvement for *h*^2^ ≥ 0.2 (Fig. 3). The improvement was not statistically significant at this value of *h*^2^ when only 100,000 samples were used in the simulation, but even in this case the number of causal variants recovered was always at least as large and typically larger when the PGS fixed effect was included in the model (Tables S10, S11). This was somewhat surprising, given that it is assumed that large sample sizes are required for accurate phenotype prediction from PGS^31^.

**Figure 3.**
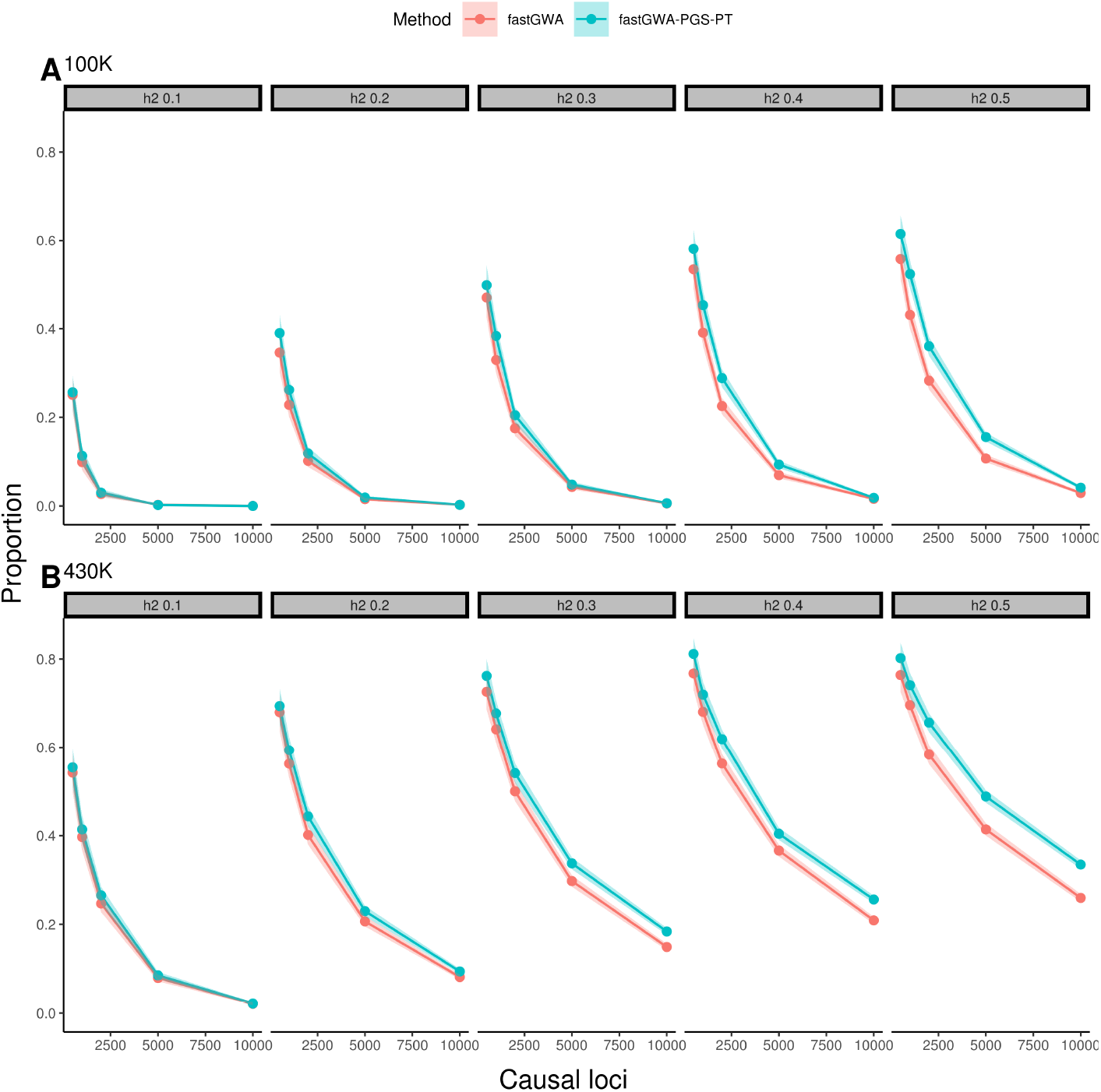
Proportion of causal variants recovered in simulations of a quantitative trait over a range of values of *h*^2^ and the number of causal loci. Simulations on the top (A) and bottom (B) panels were based on 100,000 and 430,000 randomly sampled individuals from the UK Biobank, respectively.

The improvement in power resulting from the inclusion of the PGS fixed effect increased consistently with increasing numbers of causal variants in the case of the larger sample size. This was not the case for the smaller sample size, for which the improvement decreased or was lost altogether when the number of causal variants was large (Fig. 3). This is likely due to the loss of power to detect true causal variants and to estimate their effect sizes accurately when the genetic effect is distributed over too large a number of causal variants, resulting in the inability to correct for the genetic background using the PGS. This suggests that larger sample sizes would be required for highly polygenic traits in order to obtain a benefit from using the LOCO PGS fixed effect. However, the larger sample size simulated is comparable in scale to the UK Biobank and with a sample of this size our simulations suggest that a significant improvement in power can be obtained, even for a trait with 10,000 independent causal loci. For the case control simulation (N=100,000), a more modest increase in power was observed as heritability increased, whereas the power to recover smaller effect loci decreased dramatically compared to the quantitative simulation. However, we found that for all except three simulations the inclusion of a fixed effect LOCO PGS improved the power to detect associated loci (Fig. S5, Table S12).

### Application to UK Biobank phenotypes

We assessed performance of fastGWA-PGS on real data using standing height, BMI, and heel bone mineral density (HBMD) in individuals of British ancestry (*N_height_* =395,133, *N_BMI_* =395,149 & *N_HBMD_*=229,191) from the UK Biobank. The distribution of P-values obtained from fastGWA with the LOCO PGS included was lower than that obtained using fastGWA (Fig. S6-8). At a genome-wide significance level of 5*x*10^−8^ inclusion of a LOCO PGS always increased the number of independent loci recovered, compared to fastGWA (Table 2). Across height, HBMD, and BMI, BOLT-LMM identified the largest number of independent associated loci. Including the PGS fixed effect resulted in substantial increases in the number of independent associated loci, compared to fastGWA alone for all phenotypes (Table 2, Table S13).

**Table 2.** Number of independent significant loci identified and resulting phenotype prediction model fit

| Method | Significant loci | $R^2$ full | $R^2$ pgs | Spearman's $\rho$ | Phenotype |
| --- | --- | --- | --- | --- | --- |
| BOLT-LMM | 1,804 | 0.679 | 0.170 | 0.388 | Height |
| fastGWA | 1,318 | 0.671 | 0.165 | 0.382 |  |
| fastGWA-PGS-LDpred2 | 1,717 | 0.678 | 0.176 | 0.395 |  |
| fastGWA-PGS-PT | 1,583 | 0.676 | 0.173 | 0.391 |  |
| BOLT-LMM | 583 | 0.146 | 0.134 | 0.356 | BMI |
| fastGWA | 450 | 0.143 | 0.130 | 0.351 |  |
| fastGWA-PGS-LDpred2 | 500 | 0.142 | 0.127 | 0.346 |  |
| fastGWA-PGS-PT | 493 | 0.144 | 0.130 | 0.351 |  |
| BOLT-LMM | 393 | 0.194 | 0.155 | 0.427 | HBMD |
| fastGWA | 324 | 0.189 | 0.150 | 0.418 |  |
| fastGWA-PGS-LDpred2 | 385 | 0.196 | 0.157 | 0.432 |  |
| fastGWA-PGS-PT | 365 | 0.194 | 0.155 | 0.427 |  |

One way to determine objectively whether fastGWA-PGS outperforms fastGWA on real data is to apply both methods on the key task of phenotype prediction. We partitioned the UKB data into independent training and testing datasets of European ancestry and applied, BOLT-LMM, fastGWA and fastGWA-PGS to the training data only to obtain summary statistics. We then used these summary statistics to calculate PGS scores using LDpred2 and PRSice2 (see Methods for details). For two of the three phenotypes (height and HBMD), the PGS fixed effect resulted in an increase in the correlation between the PGS and the phenotype in the test data (Table 2). In both cases the highest correlation between with the phenotype was obtained using fastGWA-PGS-LDpred2, which out-performed BOLT-LMM on this task. For the remaining phenotype (BMI), the addition of the PGS fixed effected resulted in no change or a slightly worse correlation with the phenotype in the test data. In this case the highest performance was obtained by BOLT-LMM (but at a substantial cost in terms of computational cost; Table 1). However, even in this case, we found that including only the SNPs with low P-values in the polygenic score (as implemented by the P&T method) resulted in an improvement over fastGWA (Fig. S9).

## Discussion

Omitting covariates that are associated with a response and independent of an effect of interest can result in a reduction in the efficiency of the estimation of the effect of interest^15, 16^. Complex traits are associated with the genotype of many loci across the genome, but the effects of genetic variants other than the variant being tested are often not fully modelled by GWAS methods. We evaluated a simple two-stage approach to accounting for this genetic background effect that consists of performing an initial GWAS and using the summary statistics to calculate a polygenic score and then including the polygenic score, derived from SNPs not on the same chromosome as the target SNP, as a fixed effect in a second round of association testing. Using simulated data, we found that this led to a substantial improvement in power of fastGWA, an efficient tool for biobank scale GWAS that does not fully control for genetic background effects. When we included the LOCO polygenic score as a fixed effect with fastGWA (which we refer to as fastGWA-PGS), the power exceeded that of REGENIE^30^, a recent, computationally efficient tool for GWAS that uses ridge regression to control for genetic background effects. When BOLT-LMM^8^ was used with a GRM derived from all of the simulated variants, the LOCO PGS fixed effect did not provide any boost in power (Fig. S3); however, the equivalent (or slightly improved) performance of fastGWA-PGS-LDpred2 (Fig. 1) was achieved at a much lower computational cost (Table 1). Furthermore, we note, that our simulations were favourable to BOLT-LMM because the LOCO PGS was calculated from the same set of variants that were used in the GRM of BOLT-LMM. In practice, millions of variants could be included in the LOCO PGS calculations, but the number of variants, *M*, that can be included in the GRM of BOLT-LMM is constrained by memory and compute time, both of which scale at least linearly with *M*. A further key advantage of the approach that we propose is that it is modular. Any phenotype prediction method can be used to predict the combined effect of the LOCO genetic variants on the phenotype. As methods for phenotype prediction improve, we anticipate that the performance of this approach will increase.

The increase in power using the PGS fixed effect was largest for simulated phenotypes with high heritability and a large number of causal variants (Fig. 3). In these cases the many background SNPs collectively explain a substantial proportion of the phenotypic variance and summarizing the contribution of these background SNPs to the phenotype via the LOCO PGS is likely to result in a better estimate of the effect of the target SNP and its standard error. The boost in performance derived from including the LOCO PGS as a fixed effect also depended on study sizes. For example, when the number of causal variants became large (10,000) there was no substantial boost in performance in the simulation that included 100,000 individuals, presumably because in this case the study size was not sufficient to identify and accurately estimate the effects of the causal variants. Even with this large number of causal variants the larger simulation (with 430,000 individuals) still showed a significant improvement arising from the LOCO PGS fixed effect (Fig. 3). Across all the simulation parameters we investigated, the performance of the fastGWA-PGS was never worse than fastGWA without the LOCO PGS. We also note that we calculated the pruning and thresholding LOCO PGS using SNPs that were selected based on a fixed P-value threshold. Further increases in power may be possible by optimizing the SNPs that are used to calculate the PGS separately for each omitted chromosome. This optimization was not required for LDpred2, which may help to explain why we achieved significantly better power when the LOCO PGS was calculated using this method rather than pruning and thresholding (Fig. 1).

We also applied the method to real data (standing height, heel bone mineral density (HBMD) and body mass index (BMI) in individuals of British ancestry in the UK Biobank). Consistent with the simulation results, we found more independent trait-associated loci using fastGWA-PGS-LDpred2 than with fastGWA alone for all three traits (30%, 19%, & 11% more for height, HBMD, and BMI, respectively; Table 2). Although, BOLT-LMM recovered the largest number of independent significant loci across all UK Biobank traits, this did not always translate into better correlation between a PGS calculated from the resulting summary statistics and the phenotype in the test dataset. In fact, the highest correlation was obtained by fastGWA-PGS-LDpred2 for two of the three traits. This could be explained by a higher proportion of true positives among the loci detected using the PGS-based methods or a more accurate estimate of the effects sizes by these methods, as suggested by Fig. S4. For BMI, the correlation was in fact lower between the PGS and the phenotype in the test dataset when the LOCO PGS fixed effect was used (Table2). However, even in this case a larger number of significant variants were recovered than with fastGWA. Interestingly, when only the variants with lower P-values were included for the calculation of the PGS (using the P&T method), the correlation between the PGS and the phenotype in the test dataset was higher when the PGS LOCO fixed effect was included (Fig. S9). This is consistent with a larger number of true positives (and therefore greater power) and/or better effect size estimates for the SNPs that were significantly associated with the phenotype.

The use of polygenic scores for phenotype prediction from genotype is an increasingly important application of the results of GWAS^32^. High polygenic scores can capture a substantial component of the risk of complex diseases^24, 33^ and guide interventions that can confer health benefits to individuals and reduce the stress on health systems^34^. Performing GWAS on a subset of samples and predicting on the remainder, we observed an increase in the correlation of the PGS with the phenotype when we included the LOCO PGS as a fixed effect in two out of three traits considered, consistent with improved effect size estimates (Fig. S4). Our results suggest that a modular approach that integrates advances in phenotype prediction with efficient GWAS methods can have a significant impact on the power of GWAS and that this can, in turn, lead to more accurate phenotype prediction. A recent study showed that models that allow unequal a priori contribution of SNPs to trait heritability can lead to substantial improvements in the accuracy of trait^35^. As new efficient methods emerge from these and further insights, they can be easily substituted for the calculation of the LOCO PGS fixed effect. The current fast pace of methodological innovation in phenotype prediction supports the use, at least for the time being, of the simple modular approach to modeling genetic background effects evaluated here.

## Conclusion

The tasks of detecting trait-associated variants and predicting the trait in a new sample from the summary statistics of these variants are closely intertwined. Improved performance on the trait-association task can result in more associated variants and better estimates of their effect sizes, resulting in improvement on the prediction task. On the other hand, improved methods for phenotype prediction can help to control for background genetic effects in methods that identify the trait-associated variants and their effects. The method that we have explored here consists of incorporating a LOCO PGS as a fixed-effect covariate to control for these background genetic effects; however, any method for phenotype prediction could play this role, once its application is restricted to variants that are not linked to the target SNP. We show here that incorporating the PGS as a fixed-effect covariate results in increased power to detect trait-associated variants in GWAS. The resulting trait-associated variants and effect size estimates can lead to an improvement in the PGS, as illustrated by improved performance in the task of predicting the phenotype in a test dataset.

## Methods

### Simulations

#### Genotype QC

The use of the UK Biobank Materials falls within UK Biobank’s generic Research Tissue Bank (RTB) approval from the NHS North West Research Ethics Committee, UK. The simulated genotype data was based on autosomal genotyped data from the UK Biobank. To limit the effects of population stratification only individuals reporting white British ancestry (data field 21000; code 1001; N=443,076) were included in these analyses. The genotype data for the simulation analysis was based on directly genotyped variants with minor allele frequency (MAF) greater than 0.05%. Variants with genotype missingness greater than 2% or that failed a test for Hardy-Weinberg equilibrium (HWE) at *α*= 0.0001 were excluded, resulting in a total of 664,393 genetic variants. There were 429,359 samples remaining following filtering. The sparse GRM required by fastGWA was created by setting entries corresponding to sample pairs with an estimated relatedness of less than 0.05 to 0. To account for population structure in the association studies, principal component analysis (PCA) was performed on a set of 165,684 variants LD-pruned with an *R*^2^ greater than 0.1 in a sliding window of size 500bp, sliding by 200bp. This set was also used as the basis of the BOLT-LMM analyses with the reduced GRM size (referred to as BOLT-LMM-165 in Results). All genotype QC was implemented in plink2^36^.

Based on the above genotype data, we simulated a continuous phenotype using the GCTA software suite^37^. The initial simulation consisted of 100,000 individuals, 1,000 randomly sampled causal variants and *h*^2^ = 0.5. This simulation was repeated 100 times with the 664,393 variants remaining after variant filtering for the GRM calculation. Power was calculated as the proportion of the causal variants recovered. To calculate false positive rates we first removed all SNPs within 1 Mb of the causal SNPs. Further simulations were carried out to investigate the effects of varying the number of causal SNPs, *h*^2^ and the sample size on method performance. In each case all parameters other than the ones being varied were the same as the initial simulation, and one simulation was performed per set of parameter values. The pROC R package was used to generate receiver operating characteristic (ROC) curves^38^. We applied the same simulation strategy to binary traits with two levels of disease prevalence, 0.1 & 0.3, using 1,000 causal loci with *h*^2^ = 0.5, and 100,000 samples.

#### Simulation Association tests

Association testing was performed using fastGWA, REGENIE and BOLT-LMM. To account for known sources of covariation (technical batch effects, population structure, biological effects) 10 PCs, sex, age, genotype batch and assessment centre were included as fixed-effect covariates in statistical models. For the PGS method we first performed GWAS (using fastGWA, REGENIE or BOLT-LMM) and calculated PGS scores on a Leave One Chromosome Out (LOCO) basis. This resulted in 22 sets of PGS values (one for each autosomal chromosome, calculated from the summary statistics of variants on all other autosomal chromosomes). Two PGS strategies were used in this study, pruning and thresholding (P+T), denoted with the suffix PGS-PT and LDpred2, denoted by the suffix PGS-LDpred2. The LOCO PGS-PT were calculated using PRSice2 (version 2.2.12 (2020-02-20))^28^. To decrease computation time and reduce the likelihood of over-fitting a P-value threshold of 5 × 10^−5^ was chosen, a priori, for the LOCO PGS-PT calculation. Association testing was then performed using fastGWA in a chromosome-wise manner, with the corresponding LOCO PGS included as a fixed effect. The bigsnpr R package was used to calculate the LOCO PGS-LDpred2 fixed effects^29^. To reduce computation time, 22 LOCO genotype objects containing the SNP correlations were precomputed.

### Application to the UK Biobank

#### UK biobank Association tests

The genotype selection, quality control and genetic relationship matrix were performed following the QC procedure in *Jiang et al.*^9^. The genetic relationship matrix used with fastGWA and BOLT-LMM was calculated for all European individuals (N=458,686), using a set of 556,516 lightly pruned HAPMAP3 variants (*R*^2^ greater than 0.9 in a 100 variant sliding window of size 1,000 & MAF > 0.01)^9^. Association summary statistics were generated from a set of 1.1 million HAPMAP3 variants (MAF 0.01, HWE *α*= 1 × 10^−6^ and missingness < 0.05)^9^. Principal components were calculated using a set of 34,775 variants (LD-pruned with *R*^2^ = 0.05 in a sliding window of size 1,000bp, sliding by 50bp)^39^. To identify white British samples with similar genetic backgrounds we clustered samples based on the first 6 principal components^39^, resulting in a subset of 406,319 white-British samples. Sample pairs that had a KING kinship coefficient above 0.05, with one member of the pair within the white-British group and the other in the group self-reporting as white European were removed. This left 399,135 white British and 46,406 other European samples^39, 40^. To account for known sources of phenotype and genotype variation, 10 PCs, age, sex, genotype batch and assessment centre were included as fixed-effect covariates for the BOLT-LMM and fastGWA analyses. PRSice2 and LDpred2 were used to calculate the LOCO PGS. Independent loci were identified using the clumping algorithm in plink2 (P-value threshold = 5*x*10^−9^, window size = 5Mb, and LD *R*^2^ threshold = 0.01).

#### UK Biobank phenotype prediction

To test the performance of fastGWA-PGS on the task of predicting standing height, BMI and HBMD, the UK Biobank data was partitioned into training and test datasets. The test data consisted of white British individuals with similar genetic background described above and the polygenic score predictions were tested on the remaining independent European samples. Summary statistics were generated using fastGWA, fastGWA-PGS-PT, fastGWA-PGS-LDpred2 and BOLT-LMM. We used LDpred2 and PRSice2 to predict the phenotypic values in the test set. LDpred2 requires LD correlation data and we used a pre-computed set built on the 1.1 million HAPMAP3 variants for this purpose. The model fit was assessed for each method by fitting a linear model to the values of the phenotype in the test set as a function of their predicted values, accounting for known sources of phenotypic variation, i.e sex, age, PC’s. We report both the proportion of variation explained collectively by the PGS, sex, age, the first 4 principal components and assessment centre as well as the *R*^2^ using only the PGS in the regression model.

## Data Availability

All genotype and phenotype data analyzed are available, subject to application, from the UK Biobank (application 23739). Code to implement the fastGWA-PGS method described in this work is available under MIT license from https://github.com/declan93/PGS-LMM/.

## Acknowledgements

This research has been conducted using the UK Biobank Resource under Application Number 23739. This publication has emanated from research conducted with the financial support of Science Foundation Ireland under Grant number 16/IA/4612.

## Author contributions statement

CS conceived and supervised the project and performed analyses. DB implemented the pipeline and performed analyses. DB and CS wrote the manuscript, with input from DM. DM advised on application of the method to human phenotypes. DOS performed analysis. JF provided the mathematical justification of the method.

