## Supplementary material for "Controlling for background genetic effects using polygenic scores improves the power of genome-wide association studies"

May 18, 2021

### Theory

The following supplement gives more technical arguments that conditioning on a polygenic gene score, that is constructed from SNPs on off-target chromosomes, selected for signficance of association with the outcome, improves statistical power while conserving type I error in a standard linear mixed model. To simplify arguments, we will compare the two approaches fastGWA and fastGWA-PGS. The argument will be in 3 stages. First, we derive an expression for the variance of the association estimate when the PGS is not adjusted for. Second, we derive an expression for the variance of the association estimate in the PGS adjusted model, under the assumption that the genetic and environomental residuals remain independent of the target SNP genotype conditional on the PGS - importantly, this independence condition also implies that the association parameter being estimated is the same in the models with and without adjustment for PGS. This PGS-adjusted association will be seen to have smaller variance than the corresponding estimator from the unadjusted model. Finally, we argue this independence condition (of the residuals in the PGS-adjusted model and SNP genotype) is approximately true assuming that the PGS is statistically independent of the selected

SNP. In practice, the PGS should be approximately independent of the selected SNP under the null hypothesis of no causal association at the target SNP, since the off-target SNPs that constitute the polygenic score are selected independently and are on differing chromosomes (that is they are not in LD with the target SNP and there is no-collider bias between the target SNPs and off-target SNPs since the null hypothesis is true). This proves the conservation of type I error. Under the alternative hypothesis that the target SNP has a causal association with the outcome, collider bias might result in some correlation between the PGS and target SNP genotype; however, the extent of this correlation is likely extremely weak when there are a large number of variants that are associated with the trait in question, and unlikely to invalidate the following argument.

We first list the assumptions and notation we will use for the remainder of the argument.

### Assumptions

- Let X correspond to the standardized SNP genotype at a particular location
- Without loss of generality, assume that $Var(X) = 1$ and $E(X) = 0$ (that is if $X^*$ is the original genotype data, $X = (X^* - E(X^*))/SD(X^*)$
- Similarly, the outcome $Y$ is standardized, so that $E(Y) = 0$ and $Var(Y) = 1$
- Data collected on outcome, $Y$, target SNP $X$, and offtarget genetic SNPs, $G_1, ..., G_K$ for samples $i = 1 \ldots N$
- The estimated LOCO polygenic score, $\hat{P} = \sum_{k \in \hat{S}} \hat{\beta}_k G_k$, constructed over SNPs in the selection set $\hat{S}$. Again SNP variables $G_k$ for $k \in S$ are standardized to have mean 0, variance 1. By construction, $\hat{P}$ has expected value 0. We assume that $\hat{\beta}_k$ are scaled so that the empirical variance of $\hat{P}$ over samples $i \leq N$ is 1.
- Finally, we consider the LOCO polygenic score $P$ that corresponds to SNPs in $S$ but weighted according to their "true" associations $\beta_k$, $P = \sum_{k \in \hat{S}} \beta_k G_k$

- Subscript notation. $i$ and $j$ refer to individuals $i, j \leq N$; $k \leq K$ refers to genetic location

### Variance of $\hat{\beta}$ in fastGWA model

The fastGWA model takes the form:

$$Y = \beta X + g^{(0)} + \varepsilon^{(0)} \tag{1}$$

where $Var(\varepsilon_1^{(0)}, ..., \varepsilon_N^{(0)}) = \sigma_0^2 I$ and $Var(\mathbf{g^{(0)}}) = Var(g_1^{(0)}, ..., g_N^{(0)}) = \Pi\tau_0^2$, where the family matrix $\Pi$ is assumed known (or can be estimated using the original genotypes). The overall variance matrix of $Var(\mathbf{Y}) = (Y_1, ..., Y_N)$ in (1) accounting for both the environmental variance and genetic random effect is $V = \sigma_0^2 I + \Pi\tau_0^2$ Assuming consistent REML estimates,$\hat{\tau}_0$ and $\hat{\sigma}_0$, of $\tau_0$ and $\sigma_0$, estimated by fastGWA, fastGWA estimates $\beta$ by genalized least squares:

$$\hat{\beta} = \mathbf{X}^t \hat{V}^{-1} \mathbf{Y}$$

Since, $\hat{\beta}$ is computed using generalized least squares, it is easily shown that:

$$Var(\hat{\beta}) = (\mathbf{X}^t \hat{V}^{-1} \mathbf{X})^{-1}$$

with $\mathbf{X}$ being the vector of the target SNP over $i = 1, ..., N$

Henceforth, we will assume that estimation error in the estimated variance components: $\hat{\sigma}_0$ and $\hat{\tau}_0$ is negligible, so can effectively leave out the hat-notation when referring to variance components.

To examine the effect of the extent of family correlation structure on $Var(\hat{\beta})$ in a simplistic setting, we will assume that $\Pi$ has a compound symmetry structure (implying that all individuals are equally related. That is

$$\Pi = \rho \mathbf{J} + (1-\rho)\mathbf{I}$$

where $J$ is the $N \times N$ matrix of 1's. That is $\Pi$ has elements $-1 \leq \rho \leq 1$ on its off diagonals and 1 on its diagonals. It follows that the matrix $V$ has also a compound symmetry form:

$$V = \rho \tau_0^2 \mathbf{J} + ((1-\rho)\tau_0^2 + \sigma_0^2)\mathbf{I}$$

The inverse of $V$ (if it exists) can be calculated analytically and is equal to:

$$V^{-1} = \mathbf{I}/((1-\rho)\tau_0^2 + \sigma_0^2) - \mathbf{J}\frac{\rho \tau_0^2}{((1-\rho)\tau_0^2 + \sigma_0^2)((1-\rho)\tau_0^2 + \sigma_0^2 + N\rho\tau_0^2)}$$

It follows that:

$$Var(\hat{\beta}) = (\mathbf{X}^t \hat{V}^{-1}\mathbf{X})^{-1} = [\frac{\sum_{i \leq N} X_i^2}{(1-\rho)\tau_0^2 + \sigma_0^2} - \frac{\sum_{i,j \leq N} X_i X_j \rho \tau_0^2}{((1-\rho)\tau_0^2 + \sigma_0^2)((1-\rho)\tau_0^2 + \sigma_0^2 + N\rho\tau_0^2)}]^{-1}$$

Now, noting that $E(X_i^2)$ =1 and assuming that $E(X_iX_j) = \rho$, the genetic correlation, for large $N$ one can show that the above is approximately equal to

$$Var(\hat{\beta}) = \frac{\sigma_0^2 + (1-\rho)\tau_0^2}{N(1-\rho)} \quad (2)$$

indicating that $Var(\hat{\beta})$ is smallest when fastGWA is run on unrelated individuals, that is where $\rho = 0$. From this, we see that the inclusion of a genetic-random effect (with a particular correlation matrix) in fastGWA does little to increase power (although the association estimate will be slightly more efficient than the corresponding estimate from a regression not taking into account family structure when $\rho \neq 0$. The goal in Fast-GWA is instead to properly incorporate family structure in the estimation of $Var(\hat{\beta})$. In particular, related-ness in the GWAS reduces the power of finding associated SNPs (which is indicated in that $Var(\hat{\beta})$ is a increasing function of $\rho$).

### Variance of $\hat{\beta}$ in fastGWA-PGS model

The fastGWA-PGS model takes the form:

$$Y = \beta X + g^{(1)} + \gamma\hat{P} + \varepsilon^{(1)} \tag{3}$$

where $\hat{P} = P + \varepsilon_P$ is the estimated polygenic risk score, assumed to be independent of $X$, and estimated in a LOCO fashion. We will later justify that the modified residual terms $\varepsilon^{(1)}$ and $g^{(1)}$, are zero mean random variables that are independent of $X$ conditional on $\hat{P}$ provided $\hat{P}$ is independent of $X$. Comparing with equation (1) we have that:

$$Var(\varepsilon^{(0)}) + Var(g^{(0)}) = Var(\varepsilon^{(1)}) + Var(g^{(1)}) + \gamma^2 \tag{4}$$

Importantly, these independence conditions imply that conditional on $\hat{P}$, $Cov(X,Y|\hat{P}) = \beta Var(X|\hat{P}) = \beta Var(X)$. Noting then that $Cov(X,Y|\hat{P})$ is constant, it must equal $Cov(X,Y)$, which implys that $\beta = Cov(X,Y)/Var(X)$. This indicates that the coefficient $\beta$ multiplying the SNP genotype is the same in (3) and (1). Note that the variances of both residual terms may be reduced due to addition of the polygenic risk score, that is $Var(\varepsilon^{(1)}) = \sigma_1^2 < Var(\varepsilon^{(0)}) = \sigma_0^2$ and $Var(g^{(1)}) = \tau_1^2 < Var(g^{(0)}) = \tau_0^2$. As vector equations we again assume that $Var(\varepsilon_1^{(0)},...,\varepsilon_N^{(1)})$= $\sigma_1^2 I$ and $Var(\mathbf{g^{(1)}})$ = $Var(g_1^{(1)},...,g_N^{(1)})$ = $\Pi\tau_1^2$. Comparing equations (1) and (3), it follows that adjustment for the polygenic score will reduce the variance of the environmental noise and genetic components in (1), by the quantities: $Corr(\hat{P},\varepsilon^{(0)})$ and $Corr(\hat{P},g^{(0)})$. Note if we instead adjusted for the "true" polygenic score, $P$, in the regression, we might reduce more of the noise in the genetic random effect but would not reduce noise in the environmental random effect.

The model can be approximately fit in 2 stages. First, we orthogonalize the outcome, Y with respect to $\hat{P}$. That is we set $Y^{(1)} = Y - Y_{\hat{P}} = Y - \hat{\gamma}\hat{P}$, where $Y_{\hat{P}}$ is the predicted outcome from a regression using $\hat{P}$. Second, we orthogonalize $X$ with respect to $\hat{P}$, that is calculate $X^{(1)} = X - X_{\hat{P}}$. Assuming $X$ is truly independent of $\hat{P}$ one would expect that $X^{(1)} \sim X$. Finally, $\beta$ is estimated by a generalized least squares fit, regressing $Y^{(1)}$ on $X^{(1)}$, in the following model

$$Y^{(1)} = \beta X^{(1)} + g^{(1)} + \varepsilon^{(1)} \tag{5}$$

where the variance matrix

$$V^{(1)} = \sigma_1^2 + \Pi\tau_1^2. \tag{6}$$

Similarly to before, $\hat{\beta} = \mathbf{X^{(1)}}^t \mathbf{V}^{(1)-1}\mathbf{Y^{(1)}}$ and the variance of $\hat{\beta}$ is

$$Var(\hat{\beta}) = [\mathbf{X}^{(1)t}\mathbf{V}^{(1)-1}\mathbf{X}^{(1)}]^{-1} \tag{7}$$

and under the circumstance that the off-diagonal elements of $\Pi$ are all equal to $\rho$, and $X^{(1)} \sim X$, this is approximately

$$Var(\hat{\beta}) = \frac{\sigma_1^2 + (1-\rho)\tau_1^2}{N(1-\rho)} \tag{8}$$

noting that $\sigma_1^2 < \sigma_0^2$ and $\tau_1^2 < \tau_0^2$ and comparing to (2) indicates the variance of $\hat{\beta}$ is reduced by adding the informative (and independent) estimated PGS to the regression. Because of near-orthogonality of $X$ and $\hat{P}$, one would not expect the absolute-size of $\hat{\beta}$ to be altered (indeed we argued previously that the $\beta$ coefficient in the two regression formulae (1) and (5) should be equal), indicating that a test based on $\hat{\beta}^2/Var(\hat{\beta})$ should have improved power.

### Justification of independence of modified residuals and SNP genotype $X$ under approximate independence of X and $\hat{P}$

As previously noted, if residuals, $\varepsilon^{(1)}$ and $g^{(1)}$ and genotype, $X$, in equation (3) are truly independent of each other, and $\varepsilon^{(1)}$ and $g^{(1)}$ are zero mean and finite variance, standard calculations as demonstrated later show that the variance calculated as (7) is asymptotically correct. In addition, the $\beta$ parameters will 'match' in equations (1) and (3), and hence the PGS adjusted model will have improved power under the alternative whilst conserving type I error under the null. The following is an argument to justfify this condition. By assumption, in equation (1), the residual terms $\varepsilon^{(0)}$ and $g^{(0)}$ are independent of the genotype vector $X$. We also have assumed that the selected polygenic score, $\hat{P}$ is statistically independent of $X$. This implies that once standardized to have mean 0, $X$ and $\hat{P}$ should be approximately orthogonal. Now, conditional on the vector of polygenic scores, $\hat{\mathbf{P}}$ Let $Y_{\hat{P}} = \hat{\gamma}\hat{\mathbf{P}}$ be the projection of the response vector $Y$ onto the vector $\hat{\mathbf{P}}$. By examining the right hand side of equation (1), and the approximate orthogonality of X and $\hat{\mathbf{P}}$, this projection is also equal to the sum of the projections of the vectors $\varepsilon^{(0)}$ and $g^{(0)}$ onto $\hat{\mathbf{P}}$, which we denote $\varepsilon^{(0)}_{\hat{P}} + \gamma^{(0)}_{\hat{P}}$. Now denoting $\varepsilon^{(1)} = \varepsilon^{(0)} - \varepsilon^{(0)}_{\hat{P}}$ and $g^{(1)} = g^{(0)} - g^{(0)}_{\hat{P}}$, we have the equation:

$$Y_i - \hat{\gamma}\hat{P}_i \approx \beta X_i + \varepsilon^{(1)} + g^{(1)} \tag{9}$$

where $\beta$ is the same coefficient as in equation (1). Noting that conditional on $\hat{\mathbf{P}}$, the vectors $\varepsilon^{(1)}$ and $g^{(1)}$ are functions of the vectors $\varepsilon^{(0)}$ and $g^{(0)}$, which are all independent of X, $\varepsilon^{(1)}$ and $g^{(1)}$ are also independent of X. In addition, $\varepsilon^{(0)}$, $g^{(0)}$ and $\hat{P}$ are 0-mean random variables by assumption. Since, as vectors $\varepsilon^{(1)}$ and $g^{(1)}$ can be viewed as the difference of a zero mean vector and a projection onto a zero mean vector they can also be viewed as zero mean vectors, which completes the argument.

### Conservation of Type I error, after adjustment for $\hat{P}$, assuming independence of $X$ and modified residuals

Under the scenario that we have sucessfully reduced residual noise by incorporating a polygenic risk score as above, the association test checks the orthogonality of the genotype vector for the SNP, X with the noise reduced outcome vector (after subtracting off the predicted outcome based on the polygenic score). Since the polygenic risk score is approximately othogonal to the SNP in question, and was constructed with no reference to the SNP, the Type I error of this test should not be affected. This follows in a straightforward way from the observations that the modified genetic and environmental residuals are independent of **X** and have 0 mean and the variance matrix listed above as we have justified above.

In more detail, sxuppose that $\beta = 0$. If $E(\hat{\beta}) = 0$ and the variance of $Var(\hat{\beta})$ is really given by (7), it follows that the test statistic: $\hat{\beta}^2/Var(\hat{\beta})$ should be approximately chi-squared with 1 degree of freedom, and p-values will be uniform as required for a valid statistical test.

First $E(\hat{\beta})$ = $E(\mathbf{X^{(1)}}^t\mathbf{V}^{(1)-1}\mathbf{Y^{(1)}})$ = $\mathbf{X^{(1)}}^t\mathbf{V}^{(1)-1}E(\mathbf{Y^{(1)}})$. Now since $\beta$=0, $E(\mathbf{Y^{(1)}})$ = $E(g^{(1)} + \varepsilon^{(1)}) = 0$ from the model.

Second, $Var(\hat{\beta})$ = $Var(\mathbf{X^{(1)}}^t\mathbf{V}^{(1)-1}\mathbf{Y^{(1)}})$ = $\mathbf{X^{(1)}}^t\mathbf{V}^{(1)-1}Var(\mathbf{Y^{(1)}})\mathbf{V}^{(1)-1}\mathbf{X^{(1)}}$. Now $Var(\mathbf{Y^{(1)}})$ = $Var(\varepsilon^{(\mathbf{1})}) + Var(\mathbf{g^{(1)}})$, which by definition is given by (6), implying that $Var(\hat{\beta})$ is indeed given by (7)

137 ## Supplementary figures & Tables

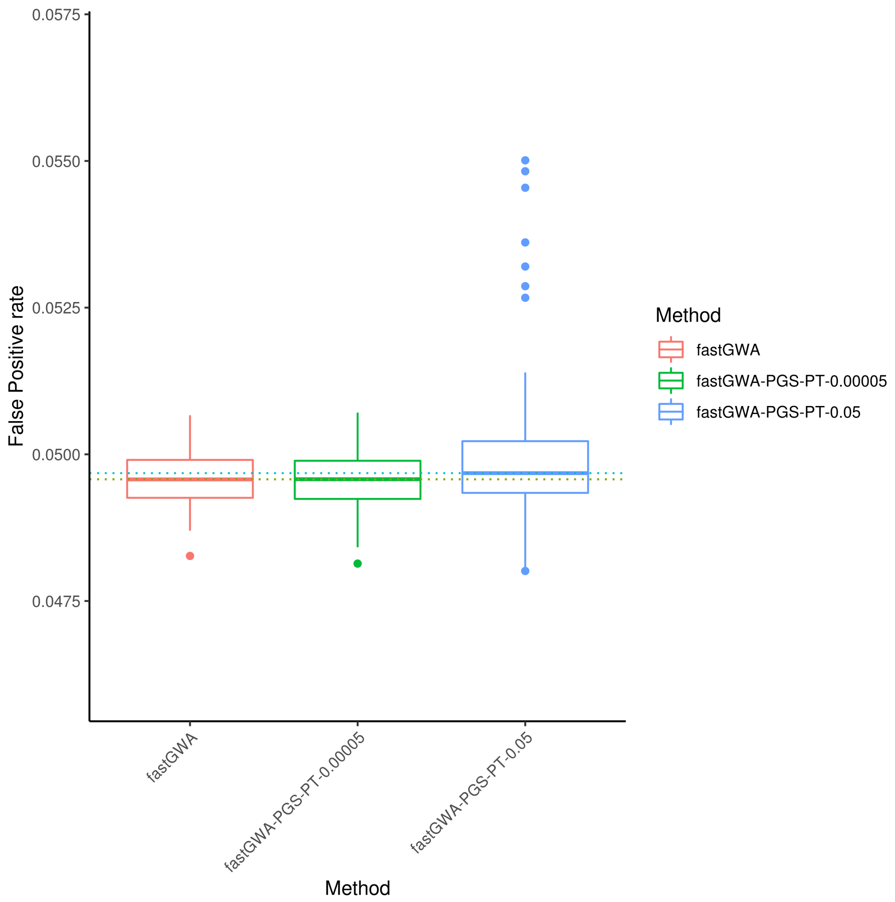


Figure S1: The false positive rate in 100 null simulations (no relationship between phenotype and genotype). The results of fastGWA-PGS-PT are shown for two P&T P-value thresholds.

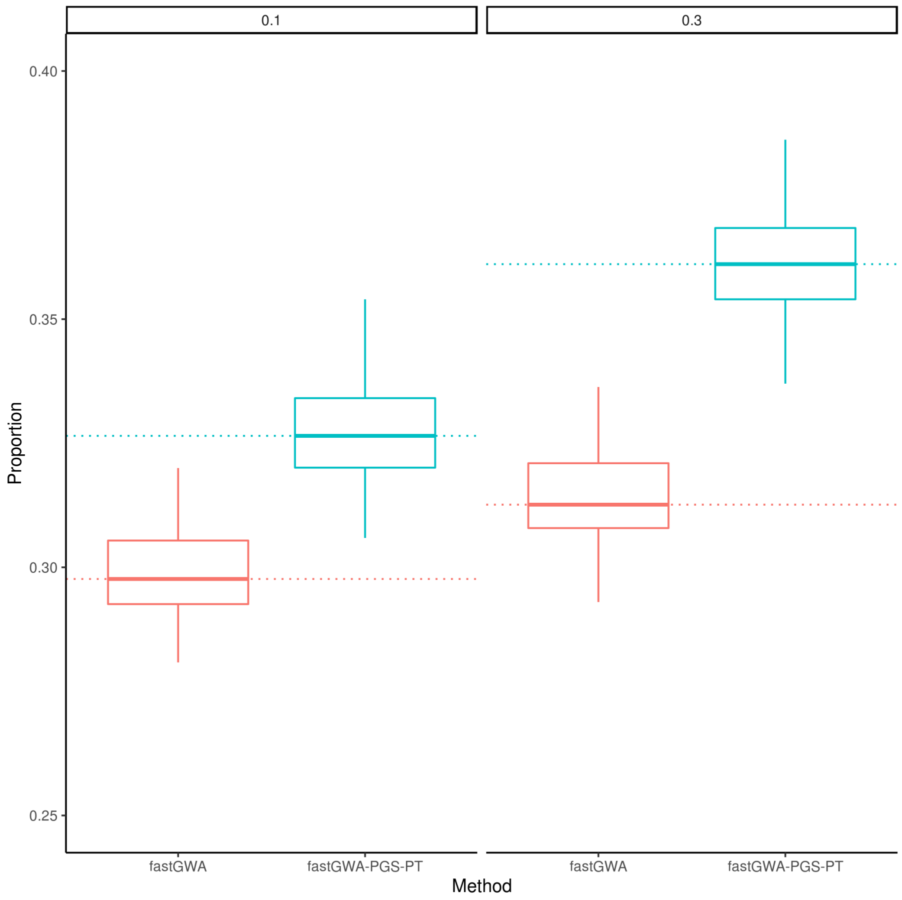


Figure S2: Proportion of causal variants recovered in 100 case-control simulations of a disease with prevalence 0.1 (left) or 0.3 (right), heritability of 0.5 and 1,000 causal variants.

Table S1: Mean proportion of causal variants recovered in 100 simulations of a quantitative trait ($h^2$=0.5, N=100,000 & 1,000 causal loci).

| Method | Mean | Change (%) relative to fastGWA |
|---:|---|---:|
| fastGWA | 0.445 | 0.00 |
| fastGWA-PGS-PT | 0.527 | 18.4 |
| fastGWA-PGS-LDPred2 | 0.561 | 25.9 |
| BOLT-LMM-165 | 0.491 | 10.3 |
| BOLT-LMM-165-PGS-PT | 0.545 | 22.4 |
| BOLT-LMM-665 | 0.558 | 25.3 |
| REGENIE | 0.481 | 8.1 |
| REGENIE-PGS-PT | 0.485 | 8.9 |

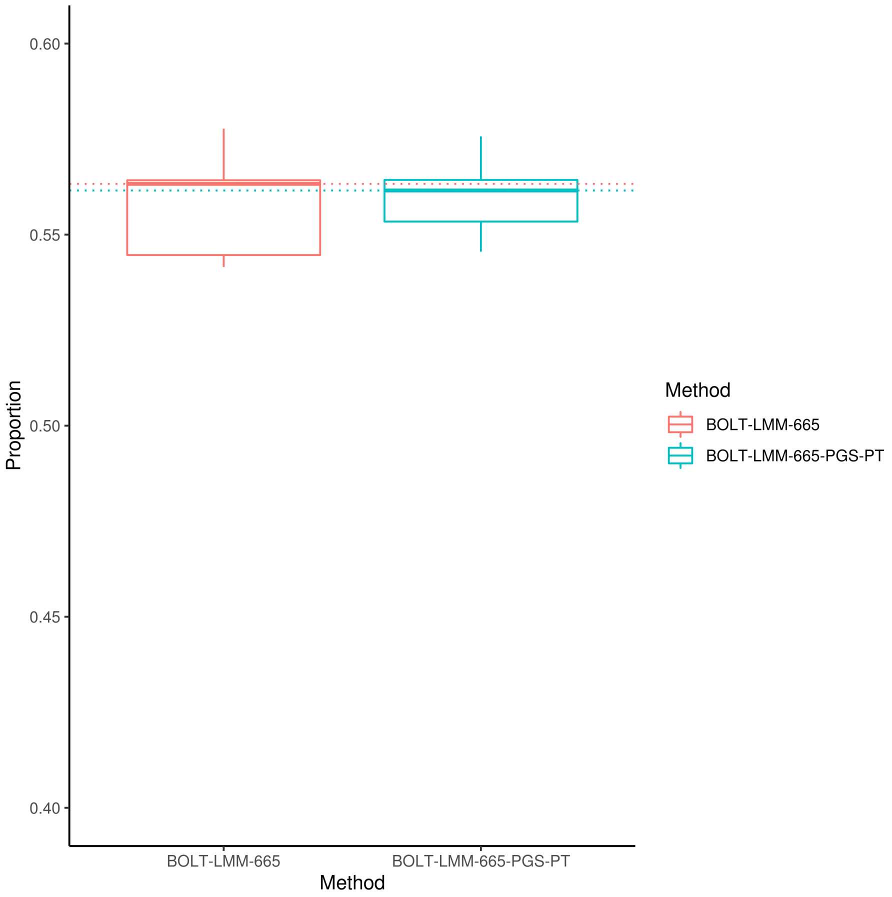


Figure S3: The effect on power of adding the LOCO PGS (based on pruning and thresholding) to BOLT-LMM with a GRM that included all variants in the simulation. The plot shows the proportion of causal variants recovered over 10 simulations.

Table S2: Paired t-tests for fastGWA vs all other methods (based on 100 simulations of a quantitative trait with $h^2$=0.5, N=100,000 & 1,000 causal loci).

| Method | Mean difference | Conf-95 | Conf+95 | P-value |
|---:|---:|---:|---:|---:|
| fastGWA-PGS-PT | 82 | 78 | 86 | 3e-32 |
| fastGWA-PGS-LDPred2 | 115 | 110 | 120 | 2.3e-36 |
| BOLT-LMM-665 | 112 | 108 | 116 | 2.3e-40 |
| BOLT-LMM-165 | 45 | 42 | 47 | 3.1e-31 |
| BOLT-LMM-165-PGS-PT | 100 | 96 | 103 | 1.4e-39 |
| REGENIE | 36 | 34 | 39 | 3e-28 |
| REGENIE-PGS-PT | 39 | 34 | 43 | 3.9e-20 |

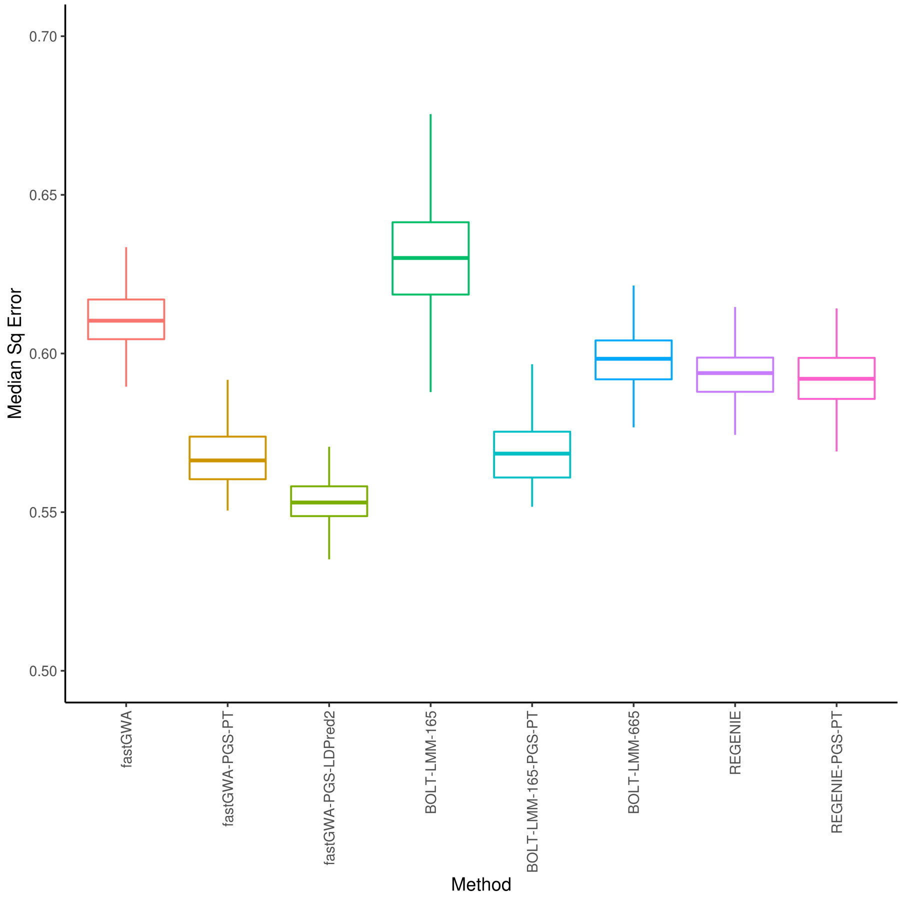


Figure S4: Median squared error of effect size estimates over 100 simulations of a quantitative trait with heritability of 0.5 and 1,000 causal variants in 100,000 individuals .

Table S3: Paired t-tests for BOLT-LMM-665 vs all other methods (based on 100 simulations of a quantitative trait with $h^2$=0.5, N=100,000, & 1,000 causal loci).

| Method | Mean difference | Conf-95 | Conf+95 | P-value |
|---:|---:|---:|---:|---:|
| fastGWA | 112 | 108 | 116 | 2.3e-40 |
| fastGWA-PGS-PT | 30 | 26 | 34 | 3.1e-18 |
| fastGWA-PGS-LDPred2 | -2.7 | -5.9 | 0.47 | 0.092 |
| BOLT-LMM-165 | 68 | 65 | 70 | 2.9e-37 |
| BOLT-LMM-165-PGS-PT | 12 | 9.9 | 15 | 1.9e-12 |
| REGENIE | 76 | 73 | 79 | 1.7e-37 |
| REGENIE-PGS-PT | 73 | 69 | 77 | 1.4e-31 |

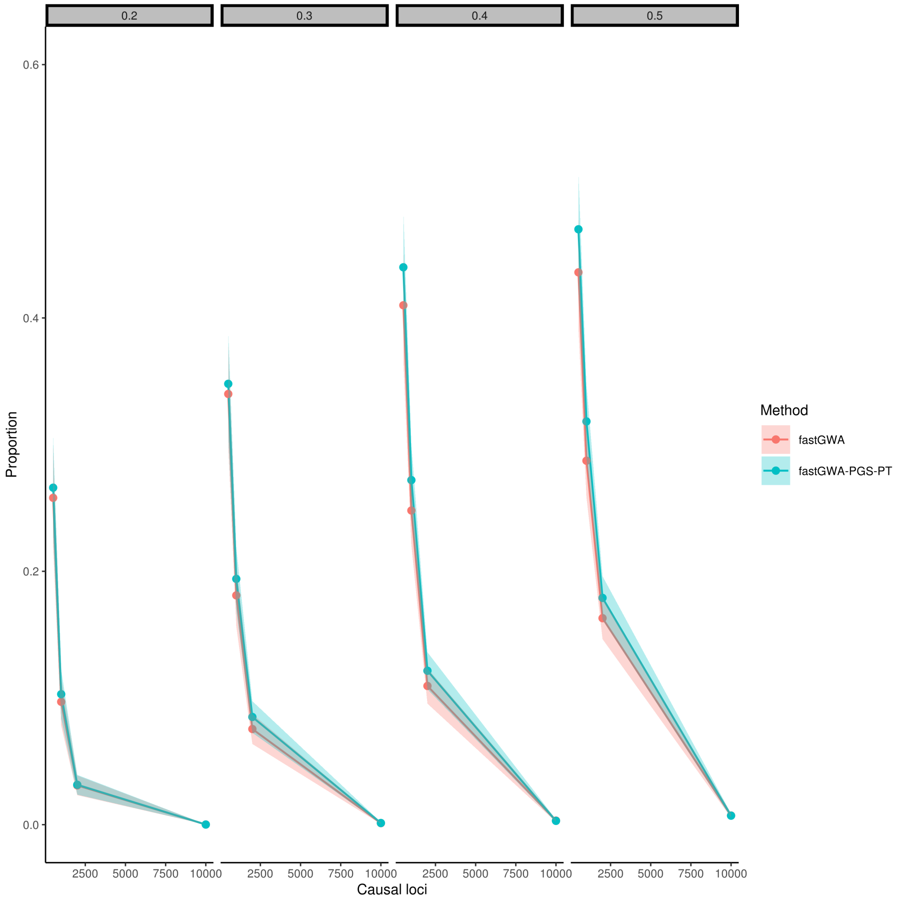


Figure S5: The proportion of causal variants recovered as a function of the number of causal variants in case-control simulations of a disease with prevalence 0.1. The plots show the results for h2 ranging from 0.2 to 0.5.

Table S4: Median proportion of recovered variants in 100 case control simulations with disease prevalence of 0.1 & 0.3 ($h^2$=0.5, N=100,000, & 1,000 causal loci).

| Method | Prevalence | Median |
|---:|:---|---:|
| fastGWA | 0.10 | 0.30 |
| fastGWA | 0.30 | 0.31 |
| fastGWA-PGS-PT | 0.10 | 0.33 |
| fastGWA-PGS-PT | 0.30 | 0.36 |

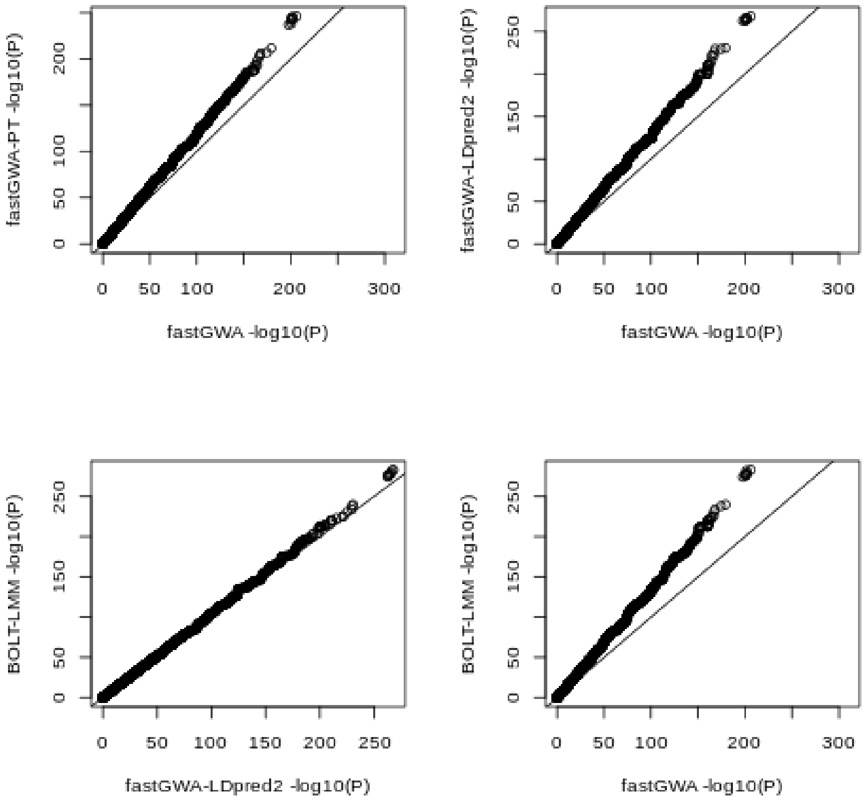


Figure S6: QQ plots comparing the distributions of the negative logarithm of the P-values obtained when different methods were applied to the height phenotype from the UK Biobank.

Table S5: Paired t-tests for 100 case control simulations with a disease prevalence of 0.1 & 0.3 ($h^2$=0.5, N=100,000 & 1,000 causal loci).

| Method | Prevalence | Mean difference | Conf-95 | Conf+95 | P-value |
|---|---|---|---|---|---|
| fastGWA-PS-PT | 0.1 | 29.3 | 28.1 | 30.6 | 2.17e-65 |
| fastGWA-PS-PT | 0.3 | 38.2 | 32.9 | 43.5 | 3.66e-25 |

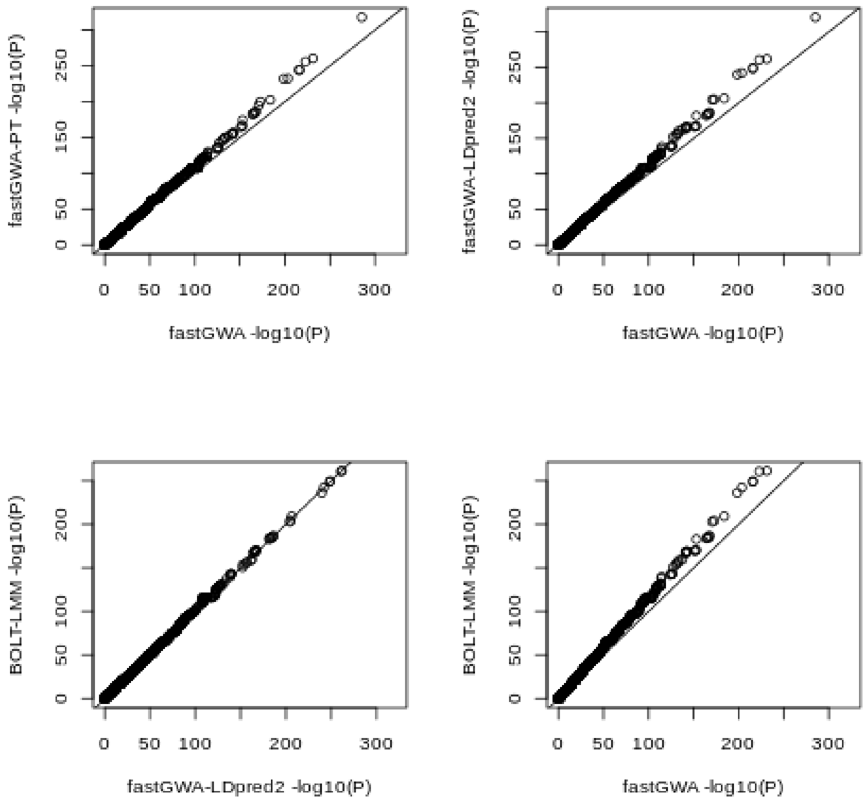


Figure S7: QQ plots comparing the distributions of the negative logarithm of the P-values obtained when different methods were applied to the heel bone mineral density (HBMD) phenotype from the UK Biobank

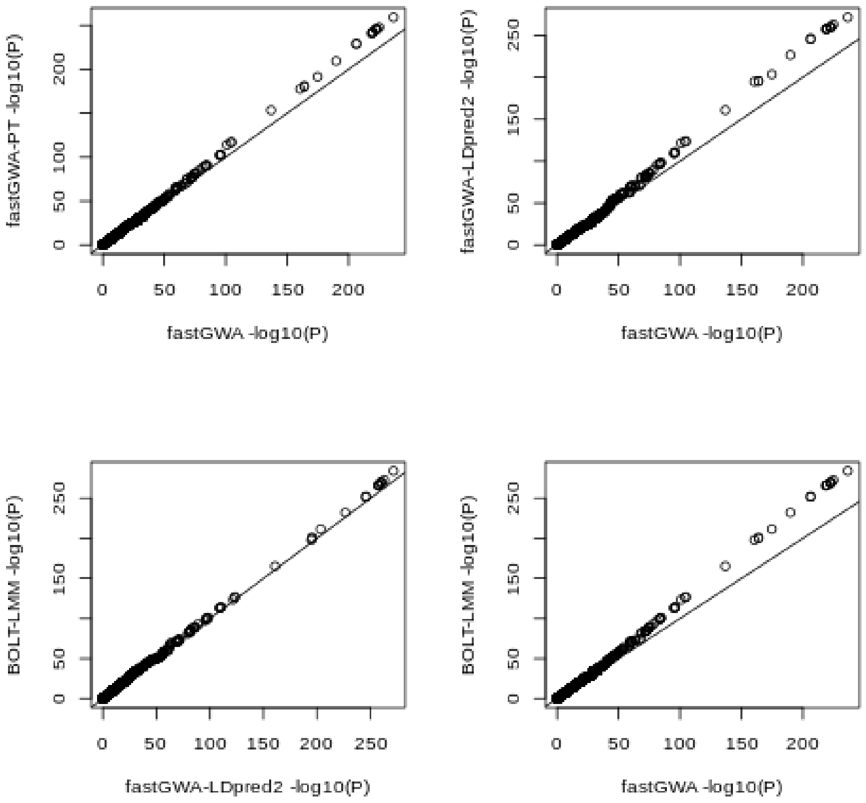


Figure S8: QQ plots comparing the distributions of the negative logarithm of the P-values obtained when different methods were applied to the body mass index (BMI) phenotype from the UK Biobank

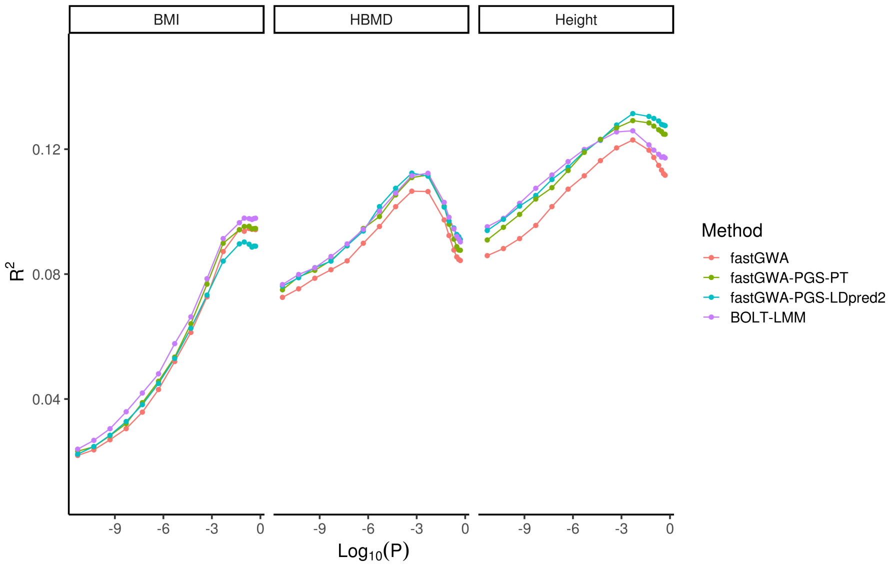


Figure S9: Proportion of phenotypic variance (in height, BMI, & HBMD) explained by polygenic scores, calculated using the P&T method, as a function of the P-value thresholds applied in the P&T method. The polygenic scores were calculated from summary statistics obtained using the methods shown.

Table S6: Maximum difference in sensitivity between methods, and the corresponding specificity at which this maximum occurs (from 100 simulations with $h^2$=0.5, N=100,000 & 1,000 causal loci).

| Method comparison | Relative increase | Max ΔSensitivity | Corresponding specificity |
|---|---|---|---|
| fastGWA-PGS-LDpred2 vs fastGWA | 0.1135 | 0.0728 | 0.9988 |
| fastGWA-PGS-LDPred2 vs BOLT-LMM-665 | 0.0016 | 0.0015 | 0.2000 |
| REGENIE vs fastGWA | 0.0315 | 0.0217 | 1.0000 |
| REGENIE-PGS-PT vs fastGWA | 0.0347 | 0.0239 | 1.0000 |
| BOLT-LMM-PGS-PT vs BOLT-LMM-165 | 0.0419 | 0.0278 | 0.9991 |
| BOLT-LMM-665 vs fastGWA | 0.1185 | 0.0766 | 0.9986 |
| fastGWA-PGS-PT vs fastGWA | 0.0847 | 0.0531 | 0.9992 |
| fastGWA-PGS-LDpred2 vs fastGWA-PGS-PT | 0.0287 | 0.0208 | 0.9966 |

Table S7: Average of the median squared error (MEDSE) of effect size estimates for causal variants across 100 simulations ($h^2$=0.5, N=100,000 & 1,000 causal loci).

| Method | Mean |
|---|---|
| fastGWA | 0.6196 |
| fastGWA-PGS-PT | 0.5756 |
| fastGWA-PGS-LDpred | 0.5612 |
| BOLT-LMM-165 | 0.6510 |
| BOLT-LMM-165-PT | 0.5764 |
| BOLT-LMM-665 | 0.6070 |
| REGENIE | 0.6032 |
| REGENIE-PT | 0.6022 |

Table S8: Paired t-tests applied to the median squared error (MEDSE) of effect size estimates for causal variants across 100 simulations, relative to fastGWA ($h^2$=0.5, N=100,000 & 1,000 causal loci).

| Method | Mean difference | Conf-95 | Conf+95 | P-value |
|---|---|---|---|---|
| fastGWA-PGS-PT | -0.044 | -0.046 | -0.042 | 3e-76 |
| fastGWA-PGS-LDPred2 | -0.058 | -0.06 | -0.057 | 5.9e-91 |
| BOLT-LMM-665 | -0.013 | -0.014 | -0.011 | 3.7e-30 |
| BOLT-LMM-165 | 0.031 | 0.015 | 0.048 | 0.00022 |
| BOLT-165-PGS-PT | -0.043 | -0.045 | -0.041 | 2.6e-71 |
| REGENIE-PGS-PT | -0.017 | -0.02 | -0.015 | 2.2e-26 |
| REGENIE | -0.016 | -0.018 | -0.015 | 1.9e-38 |

Table S9: Paired t-test of MEDSE beta estimates of 100 quantitative trait simulations relative to BOLT-LMM-165

| Method | Mean difference | Conf-95 | Conf+95 | P-value |
|---|---|---|---|---|
| fastGWA | -0.031 | -0.048 | -0.015 | 0.00022 |
| fastGWA-PGS-PT | -0.075 | -0.092 | -0.059 | 5.2e-15 |
| fastGWA-PGS-LDPred2 | -0.09 | -0.11 | -0.073 | 3e-18 |
| BOLT-LMM-665 | -0.044 | -0.061 | -0.027 | 1.1e-06 |
| BOLT-165-PGS-PT | -0.075 | -0.09 | -0.059 | 2e-15 |
| REGENIE-PGS-PT | -0.049 | -0.063 | -0.034 | 1.8e-09 |
| REGENIE | -0.048 | -0.063 | -0.033 | 1.2e-08 |

Table S10: Proportion of causal variants recovered for simulations of a quantitative trait over a range of parameter values (N=100,000; Nc = number of causal variants)

| Heritability | Method | Nc | Proportion |
|---|---|---|---|
| 0.1 | fastGWA | 500 | 0.25 |
| 0.1 | fastGWA | 1,000 | 0.10 |
| 0.1 | fastGWA | 2,000 | 0.03 |
| 0.1 | fastGWA | 5,000 | 0.00 |
| 0.1 | fastGWA | 10,000 | 0.00 |
| 0.1 | fastGWA-PGS-PT | 500 | 0.26 |
| 0.1 | fastGWA-PGS-PT | 1,000 | 0.11 |
| 0.1 | fastGWA-PGS-PT | 2,000 | 0.03 |
| 0.1 | fastGWA-PGS-PT | 5,000 | 0.00 |
| 0.1 | fastGWA-PGS-PT | 10,000 | 0.00 |
| 0.2 | fastGWA | 500 | 0.35 |
| 0.2 | fastGWA | 1,000 | 0.23 |
| 0.2 | fastGWA | 2,000 | 0.10 |
| 0.2 | fastGWA | 5,000 | 0.02 |
| 0.2 | fastGWA | 10,000 | 0.00 |
| 0.2 | fastGWA-PGS-PT | 500 | 0.39 |
| 0.2 | fastGWA-PGS-PT | 1,000 | 0.26 |
| 0.2 | fastGWA-PGS-PT | 2,000 | 0.12 |
| 0.2 | fastGWA-PGS-PT | 5,000 | 0.02 |
| 0.2 | fastGWA-PGS-PT | 10,000 | 0.00 |
| 0.3 | fastGWA | 500 | 0.47 |
| 0.3 | fastGWA | 1,000 | 0.33 |
| 0.3 | fastGWA | 2,000 | 0.18 |
| 0.3 | fastGWA | 5,000 | 0.04 |
| 0.3 | fastGWA | 10,000 | 0.01 |
| 0.3 | fastGWA-PGS-PT | 500 | 0.50 |
| 0.3 | fastGWA-PGS-PT | 1,000 | 0.38 |
| 0.3 | fastGWA-PGS-PT | 2,000 | 0.20 |
| 0.3 | fastGWA-PGS-PT | 5,000 | 0.05 |
| 0.3 | fastGWA-PGS-PT | 10,000 | 0.01 |
| 0.4 | fastGWA | 500 | 0.53 |
| 0.4 | fastGWA | 1,000 | 0.39 |
| 0.4 | fastGWA | 2,000 | 0.23 |
| 0.4 | fastGWA | 5,000 | 0.07 |
| 0.4 | fastGWA | 10,000 | 0.02 |
| 0.4 | fastGWA-PGS-PT | 500 | 0.58 |
| 0.4 | fastGWA-PGS-PT | 1,000 | 0.45 |
| 0.4 | fastGWA-PGS-PT | 2,000 | 0.29 |
| 0.4 | fastGWA-PGS-PT | 5,000 | 0.09 |
| 0.4 | fastGWA-PGS-PT | 10,000 | 0.02 |
| 0.5 | fastGWA | 500 | 0.56 |
| 0.5 | fastGWA | 1,000 | 0.43 |
| 0.5 | fastGWA | 2,000 | 0.28 |
| 0.5 | fastGWA | 5,000 | 0.11 |
| 0.5 | fastGWA | 10,000 | 0.03 |
| 0.5 | fastGWA-PGS-PT | 500 | 0.62 |
| 0.5 | fastGWA-PGS-PT | 1,000 | 0.52 |
| 0.5 | fastGWA-PGS-PT | 2,000 | 0.36 |
| 0.5 | fastGWA-PGS-PT | 5,000 | 0.16 |
| 0.5 | fastGWA-PGS-PT | 10,000 | 0.04 |

Table S11: Proportion of causal variants recovered for simulations of a quantitative trait over a range of parameter values (N=430,000; Nc = number of causal variants)

| Heritability | Method | Nc | Proportion |
|---|---|---|---|
| 0.1 | fastGWA | 500 | 0.54 |
| 0.1 | fastGWA | 1,000 | 0.40 |
| 0.1 | fastGWA | 2,000 | 0.25 |
| 0.1 | fastGWA | 5,000 | 0.08 |
| 0.1 | fastGWA | 10,000 | 0.02 |
| 0.1 | fastGWA-PGS-PT | 500 | 0.55 |
| 0.1 | fastGWA-PGS-PT | 1,000 | 0.41 |
| 0.1 | fastGWA-PGS-PT | 2,000 | 0.27 |
| 0.1 | fastGWA-PGS-PT | 5,000 | 0.08 |
| 0.1 | fastGWA-PGS-PT | 10,000 | 0.02 |
| 0.2 | fastGWA | 500 | 0.68 |
| 0.2 | fastGWA | 1,000 | 0.56 |
| 0.2 | fastGWA | 2,000 | 0.40 |
| 0.2 | fastGWA | 5,000 | 0.21 |
| 0.2 | fastGWA | 10,000 | 0.08 |
| 0.2 | fastGWA-PGS-PT | 500 | 0.69 |
| 0.2 | fastGWA-PGS-PT | 1,000 | 0.59 |
| 0.2 | fastGWA-PGS-PT | 2,000 | 0.44 |
| 0.2 | fastGWA-PGS-PT | 5,000 | 0.23 |
| 0.2 | fastGWA-PGS-PT | 10,000 | 0.09 |
| 0.3 | fastGWA | 500 | 0.73 |
| 0.3 | fastGWA | 1,000 | 0.64 |
| 0.3 | fastGWA | 2,000 | 0.50 |
| 0.3 | fastGWA | 5,000 | 0.30 |
| 0.3 | fastGWA | 10,000 | 0.15 |
| 0.3 | fastGWA-PGS-PT | 500 | 0.76 |
| 0.3 | fastGWA-PGS-PT | 1,000 | 0.68 |
| 0.3 | fastGWA-PGS-PT | 2,000 | 0.54 |
| 0.3 | fastGWA-PGS-PT | 5,000 | 0.34 |
| 0.3 | fastGWA-PGS-PT | 10,000 | 0.18 |
| 0.4 | fastGWA | 500 | 0.77 |
| 0.4 | fastGWA | 1,000 | 0.68 |
| 0.4 | fastGWA | 2,000 | 0.56 |
| 0.4 | fastGWA | 5,000 | 0.37 |
| 0.4 | fastGWA | 10,000 | 0.21 |
| 0.4 | fastGWA-PGS-PT | 500 | 0.81 |
| 0.4 | fastGWA-PGS-PT | 1,000 | 0.72 |
| 0.4 | fastGWA-PGS-PT | 2,000 | 0.62 |
| 0.4 | fastGWA-PGS-PT | 5,000 | 0.40 |
| 0.4 | fastGWA-PGS-PT | 10,000 | 0.26 |
| 0.5 | fastGWA | 500 | 0.76 |
| 0.5 | fastGWA | 1,000 | 0.70 |
| 0.5 | fastGWA | 2,000 | 0.58 |
| 0.5 | fastGWA | 5,000 | 0.41 |
| 0.5 | fastGWA | 10,000 | 0.26 |
| 0.5 | fastGWA-PGS-PT | 500 | 0.80 |
| 0.5 | fastGWA-PGS-PT | 1,000 | 0.74 |
| 0.5 | fastGWA-PGS-PT | 2,000 | 0.66 |
| 0.5 | fastGWA-PGS-PT | 5,000 | 0.49 |
| 0.5 | fastGWA-PGS-PT | 10,000 | 0.33 |

Table S12: Proportion of causal variants recovered for simulations of a binary trait over a range of parameter values (N=100,000; disease prevalence = 0.1; Nc = number of causal variants)

| Heritability | Nc | Method | Proportion |
|---|---|---|---|
| 0.2 | 10,000 | fastGWA | 0.0001 |
| 0.2 | 10,000 | fastGWA-PGS-PT | 0.0001 |
| 0.2 | 1,000 | fastGWA | 0.0970 |
| 0.2 | 1,000 | fastGWA-PGS-PT | 0.1030 |
| 0.2 | 2,000 | fastGWA | 0.0310 |
| 0.2 | 2,000 | fastGWA-PGS-PT | 0.0315 |
| 0.2 | 500 | fastGWA | 0.2580 |
| 0.2 | 500 | fastGWA-PGS-PT | 0.2660 |
| 0.3 | 10,000 | fastGWA | 0.0012 |
| 0.3 | 10,000 | fastGWA-PGS-PT | 0.0013 |
| 0.3 | 1,000 | fastGWA | 0.1810 |
| 0.3 | 1,000 | fastGWA-PGS-PT | 0.1940 |
| 0.3 | 2,000 | fastGWA | 0.0755 |
| 0.3 | 2,000 | fastGWA-PGS-PT | 0.0850 |
| 0.3 | 500 | fastGWA | 0.3400 |
| 0.3 | 500 | fastGWA-PGS-PT | 0.3480 |
| 0.4 | 10,000 | fastGWA | 0.0032 |
| 0.4 | 10,000 | fastGWA-PGS-PT | 0.0030 |
| 0.4 | 1,000 | fastGWA | 0.2480 |
| 0.4 | 1,000 | fastGWA-PGS-PT | 0.2720 |
| 0.4 | 2,000 | fastGWA | 0.1095 |
| 0.4 | 2,000 | fastGWA-PGS-PT | 0.1215 |
| 0.4 | 500 | fastGWA | 0.4100 |
| 0.4 | 500 | fastGWA-PGS-PT | 0.4400 |
| 0.5 | 10,000 | fastGWA | 0.0072 |
| 0.5 | 10,000 | fastGWA-PGS-PT | 0.0071 |
| 0.5 | 1,000 | fastGWA | 0.2873 |
| 0.5 | 1,000 | fastGWA-PGS-PT | 0.3183 |
| 0.5 | 2,000 | fastGWA | 0.1630 |
| 0.5 | 2,000 | fastGWA-PGS-PT | 0.1790 |
| 0.5 | 500 | fastGWA | 0.4360 |
| 0.5 | 500 | fastGWA-PGS-PT | 0.4700 |

Table S13: Two-sample tests of equality of proportions applied to the proportions of significant loci identified using the method shown, compared to fastGWA. The results shown are for the three UK Biobank quantitative traits analyzed. Prop 1 and Prop 2 show the proportions of significant loci for the method on the row and for fastGWA, respectively. Conf-95 and Conf+95 show the low and upper 95% confidence interval for the difference in these proportions.

| Method | Phenotype | P-value | X-sq | Prop 1 | Prop 2 | Conf-95 | Conf+95 |
|---|---|---|---|---|---|---|---|
| BOLT-LMM | BMI | 3.133e-05 | 17.3357 | 0.0159 | 0.0123 | 0.0019 | 0.0054 |
| fastGWA-PGS-LDpred2 | | 0.1083 | 2.5795 | 0.0136 | 0.0123 | -0.0003 | 0.0030 |
| fastGWA-PGS-PT | | 0.1563 | 2.0093 | 0.0135 | 0.0123 | -0.0005 | 0.0029 |
| BOLT-LMM | HBMD | 0.009114 | 6.8004 | 0.0106 | 0.0087 | 0.0005 | 0.0033 |
| fastGWA-PGS-LDpred2 | | 0.02029 | 5.3864 | 0.0104 | 0.0087 | 0.0003 | 0.0031 |
| fastGWA-PGS-PT | | 0.1066 | 2.6034 | 0.0098 | 0.0087 | -0.0002 | 0.0026 |
| BOLT-LMM | Height | 5.458e-19 | 79.2553 | 0.0493 | 0.0360 | 0.0104 | 0.0163 |
| fastGWA-PGS-LDpred2 | | 1.953e-13 | 54.0515 | 0.0468 | 0.0360 | 0.0079 | 0.0138 |
| fastGWA-PGS-PT | | 1.166e-06 | 23.6328 | 0.0430 | 0.0360 | 0.0042 | 0.0099 |
